# Integrated forward and reverse degradomics uncovers the proteolytic landscape of aortic aneurysms and the roles of MMP9 and mast cell chymase

**DOI:** 10.1101/2024.06.26.600914

**Authors:** Sumit Bhutada, Daniel R. Martin, Frank Cikach, Emidio Germano da Silva, Belinda B. Willard, Bhama Ramkhelawon, Mina K. Chung, Satakshi Dahal, Anand Ramamurthi, Jayadev P. Joshi, Daniel Blankenburg, John Barnard, Eugene H. Blackstone, Eric E. Roselli, Suneel S. Apte

## Abstract

**Background:** Dysregulated proteolysis is implicated in thoracic (TAA) and abdominal aortic aneurysm (AAA) pathogenesis, but the proteolytic landscapes (degradomes) of aneurysmal and normal aorta, and contributions of individual proteases remain undefined. Here, a proteome-wide approach was used to uncover TAA and AAA degradomes, compare them quantitatively and define the specific role in aortic remodeling of two proteases consistently identified in the aneurysms, mast cell chymase (CMA1) and matrix metalloprotease 9 (MMP9).

**Methods:** The mass spectrometry-based N-terminomics strategy <u>T</u>erminal <u>A</u>mine Isotopic <u>L</u>abeling of <u>S</u>ubstrates (TAILS) was applied to Marfan syndrome TAAs (n=5), AAAs (n=16) and corresponding non-diseased aorta (TAs, n=4, and AAs, n=8) as a forward degradomics application, i.e., to define substrate and protease degradomes, and 8-plex iTRAQ-TAILS was used for quantitative comparison. Cleavage sites of CMA1 and MMP9 were sought by reverse degradomics, i.e., digestion of aortic proteins with these proteases, followed by 6-plex iTRAQ-TAILS. CMA1 and MMP9 proteolysis of biglycan was investigated using <u>A</u>mino-<u>T</u>erminal <u>O</u>riented <u>M</u>ass spectrometry of <u>S</u>ubstrates (ATOMS).

**Results:** We experimentally annotated 16,923 proteolytically derived peptides (substrate degradome) and 90 proteases (protease degradome) in the aorta. Quantitative substrate degradome comparisons identified specific differentially modulated pathways and networks in TAA and AAA. Reverse degradomics elucidated > 300 CMA1 and MMP9 substrate cleavage sites, of which, many, including orthogonally validated biglycan cleavage, occurred in the disease degradomes.

**Conclusions:** Unbiased, proteome-wide forward degradomics of the aortic wall from TAA, AAA and non-diseased tissue generated the first systems biology view of vascular wall breakdown and public resource for the hitherto occult proteolytic landscape, demonstrating widespread extracellular matrix remodeling. The findings provide insights on aortic aneurysm pathways and potential disease biomarkers. Mapping of specific contributions of CMA1 and MMP9 on the aortic forward substrate degradome using reverse degradomics provides a strategy for defining the activities of all proteases involved in aortic disease.

## Introduction

The aorta, the prototypic elastic artery and largest blood vessel, is susceptible to numerous disorders that ultimately affect its structural integrity, leading to aneurysms. TAAs and AAAs have distinct epidemiologic associations and histopathologic characteristics. The ascending aorta, affected in Stanford type A dissections, is typically most affected by mutations in genes encoding the smooth muscle cell-extracellular matrix (ECM) continuum, which serves as an integrated mechanosensory and adaptive mechanism in aortic wall homeostasis [1]. A histopathologic hallmark of TAAs is the triad of elastic fiber fragmentation, proteoglycan accumulation, and smooth muscle cell (SMC) loss termed aortic medial degeneration [2]. AAAs typically lack this specific histopathology and are frequently associated with mural thrombus [3]. Aortic aneurysms often remain asymptomatic until catastrophic failure, typically by TAA dissection or AAA rupture, consequent to structural failure resulting from extreme remodeling [4]. Developing biomarkers for detection of silent aneurysms and monitoring of aneurysm growth are therefore major clinical needs but remain unmet.

Despite distinct etiologies, TAAs and AAAs result from extensive vascular cell and extracellular matrix (ECM) remodeling. Dysregulated aortic ECM remodeling, exemplified by elastic fiber fragmentation, alters aortic wall material properties, contributing to progressive dilatation and failure [3]. Because ECM remodeling requires proteolysis of its components, proteases have a major, yet poorly resolved role in aortic aneurysms [5–8]. The human genome encodes over 600 proteases, including numerous secreted and cell-surface proteases that attack ECM and can disrupt cell-matrix interactions [9]. As in other tissues, proteolysis may be widespread and underestimated in the aortic wall. For example, one study showed that nearly 50% of proteins in skin were cleaved to stable fragments [10] and recent analysis of osteoarthritic cartilage highlighted pervasive proteolysis [11, 12]. Shotgun proteomics analysis, where all proteins and their fragments are trypsinized into small peptides prior to mass spectrometry, and other proteomics methods typically report protein abundance and provide little information of the scale of proteolysis in aortic disease.

Since proteolysis is irreversible, it impacts tissue structure and function in myriad ways, such as by activating and inactivating proteins, releasing bound ligands, altering adhesive sites, and generating bioactive fragments. Within the aortic wall, proteolysis can destroy desirable material properties of key structural components such as collagens, elastin and proteoglycans. Phenotype modulation of vascular smooth muscle cells (SMCs), fibroblasts and other cells may be promoted by proteolytic fragments or dysregulated cell adhesion [13, 14]. Among ECM-degrading proteases, MMPs, plasminogen activators, ADAMTS proteases and mast cell chymase have recognized significance in aneurysms, with MMPs identified as drug targets [15]. Because MMP9 levels are high in both TAAs and AAAs [16], serum MMP9 was previously suggested as a diagnostic marker for aortic aneurysms [17]. Mast cells release mast cell chymase (CMA1), tryptase and carboxypeptidase A in the aorta [18, 19]. In the elastase perfusion-induced AAA mouse model, inactivation of the CMA1 homolog *Mcpt4^-/-^* reduced the number of lesional inflammatory cells and apoptosis to have an overall mitigating effect [18]. Human chymases can activate MMPs and may have many other potential substrates [18]. Despite relevance of MMP9, CMA1 and other proteases to aortic disease, their activities, i.e., their mechanisms in aneurysms, remain poorly defined.

This study had four objectives, **i**. To define the proteolytic landscape of aneurysms affecting the ascending thoracic aorta and abdominal aorta as well as of the healthy aortic wall from these regions by identifying cleaved peptide bonds, **ii**. To seek quantitative proteolytic differences associated with the distinct pathologies of TAA and AAA, **iii**. To identify proteases present in the aortic wall and **iv**. To specify the proteolytic contributions of CMA1 and MMP9 to the aortic degradome. Addressing these objectives has generated new mechanistic insights on vascular wall remodeling and resources for understanding proteolysis in human aortic disease that will ultimately facilitate monitoring of aneurysm growth and treatment.

## METHODS

A description of all experimental procedures, information on antibodies and primers and a major resources table can be found in the Supplemental Material. The data supporting the conclusions of this study are available from the corresponding authors upon request and proteomics data are available from ProteomeXChange.

## Results

### The substrate degradomes of aneurysm and non-diseased human aorta

We designed an unbiased forward degradomics strategy, in which proteolytic cleavages and proteases were identified de novo in aortic tissues (Figure 1A) to identify cleaved peptide bonds in TAA, AAA, control thoracic aorta (TA) and control abdominal aorta (AA) tissue cohorts. Two-step tissue extraction improved degradome coverage, with 47% increased yield in the second extraction step (Figure S1A). A qualitative (single-channel dimethyl TAILS) and quantitative (8-plex iTRAQ-TAILS) analysis with extraction of informative peptides from TAILS as well as pre-TAILS datasets further expanded the degradome (Figure 1B).

**Figure 1.**
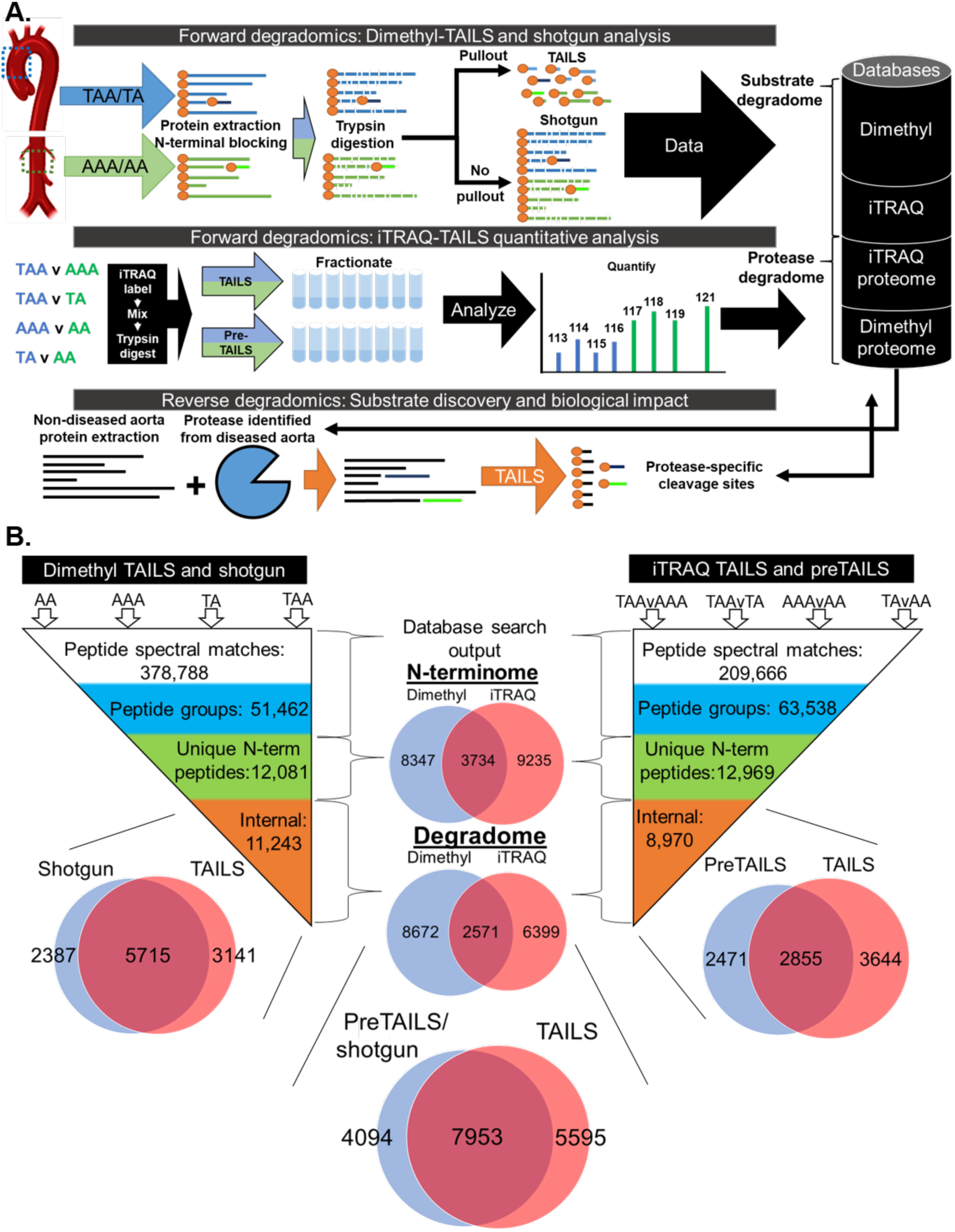
Integration of forward and reverse degradomics to generate an aortic degradome. A. Human thoracic aortic aneurysm (TAA), abdominal aortic aneurysm (AAA) and donor thoracic (TA) and abdominal aorta (AA) were analyzed using dimethyl-TAILS and 8-plex iTRAQ-TAILS to determine aortic substrate and protease degradomes. For reverse degradomics, aortic protein extracts were digested with proteases consistently identified in aneurysms (CMA1 and MMP9) followed by dimethyl-TAILS to identify substrates and cleavage sites, for matching to the aortic substrate degradome. B. Hierarchical workflow of data analysis to obtain first the N-terminome and then the degradome independently from dimethyl- and iTRAQ-TAILS analyses.

The combined tissue cohorts identified a total of 21,316 proteolytically cleaved peptides, of which 12,081 and 12,969 N-terminally blocked peptides were identified from dimethyl and iTRAQ labeled datasets respectively. Only 18% (3,734) overlapped (Figure 1B), possibly because each labeling method requires different LC-MS/MS collisional methods favoring detection of distinct peptide spectra. These peptides originated from 2,342 (dimethyl) and 2,317 proteins (iTRAQ) and among them, a large proportion were peptides representing positionally internal N-termini, comprising the aortic degradome (Figures 1B, 2A, B). By sorting the degradome into non-independent protein categories, we determined cleavages of 160 ECM proteins, of which 20 were proteoglycans, having 6,428 and 878 cleavage sites, respectively with a greater proportion of cleavage sites/protein than the other groups (Figure 2C). The number of cleaved proteins and internal peptides was consistently higher in AAAs and TAAs than in respective control cohorts with the highest per-molecule number of cleavage sites in proteoglycans and ECM (Figure 2D). These data suggested that ECM, and especially ECM proteoglycans, were disproportionately targeted by proteases in the aorta (Figure 2C, D).

**Figure 2.**
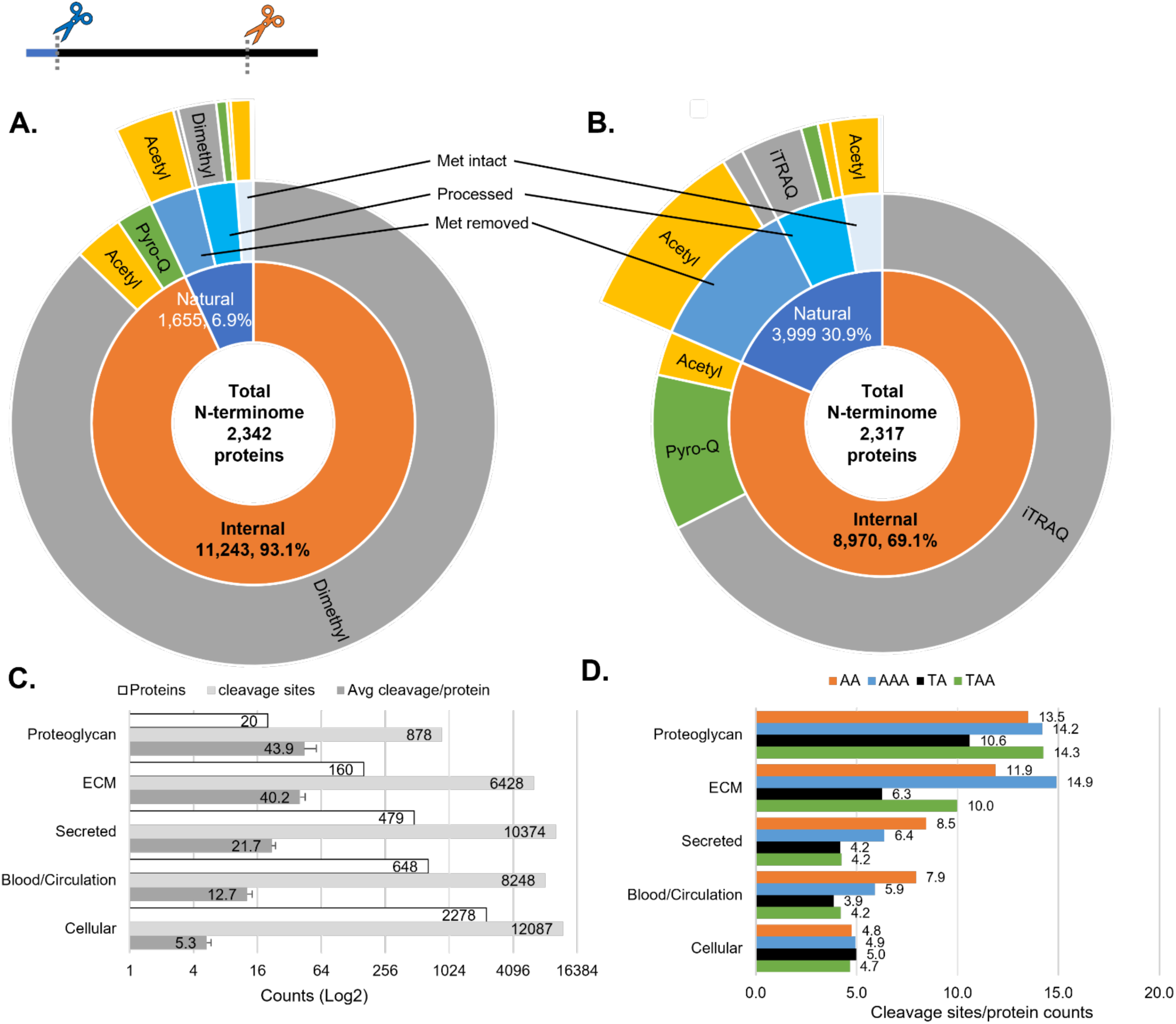
The aortic substrate N-terminome and degradome show natural N-terminal modifications and cleavages in all tissue components. A,B. Sunburst plots of peptides with blocked N-termini classified as either internal (orange) or natural (blue) based on position in the protein sequence from the dimethyl (A) or iTRAQ (B) analysis. C. Number of cleaved proteins, peptides indicative of cleavage sites and cleavage frequency for each indicated protein category. Error bars represent the variation between aortic cohorts. D. Number of peptides indicative of cleavage sites in each indicated protein category by aorta cohort.

The most numerous cleaved peptides were identified by dimethyl-TAILS in the AAA degradome with 8,649 total and 5,423 identified exclusively in this cohort. In four-way overlap analysis, TAAs and AAAs shared the most internal peptides (1,967) of any two groups (Figure 3A). Most peptides with blocked N-termini were positionally internal (Figure 3B). The number of cleavages found in the top 10 abundant proteins were defined for each cohort (Figures 3C, S1B). The most abundant protein was also the most cleaved protein in TAs (vimentin, 50 internal peptides), the second most abundant protein was the most cleaved protein in AAAs (COL6A3, 274 internal peptides) and AAs (actin, 109 internal peptides), and the most abundant protein was the second most cleaved protein in TAAs (vimentin, 81 internal peptides) (Figures 3C, S1B). A Pearson correlation plot of protein abundance and internal peptides identified per protein showed a moderate (0.3-0.5) to high (r>0.5) correlation of the two datasets in all cohorts (Figure S1C).

**Figure 3.**
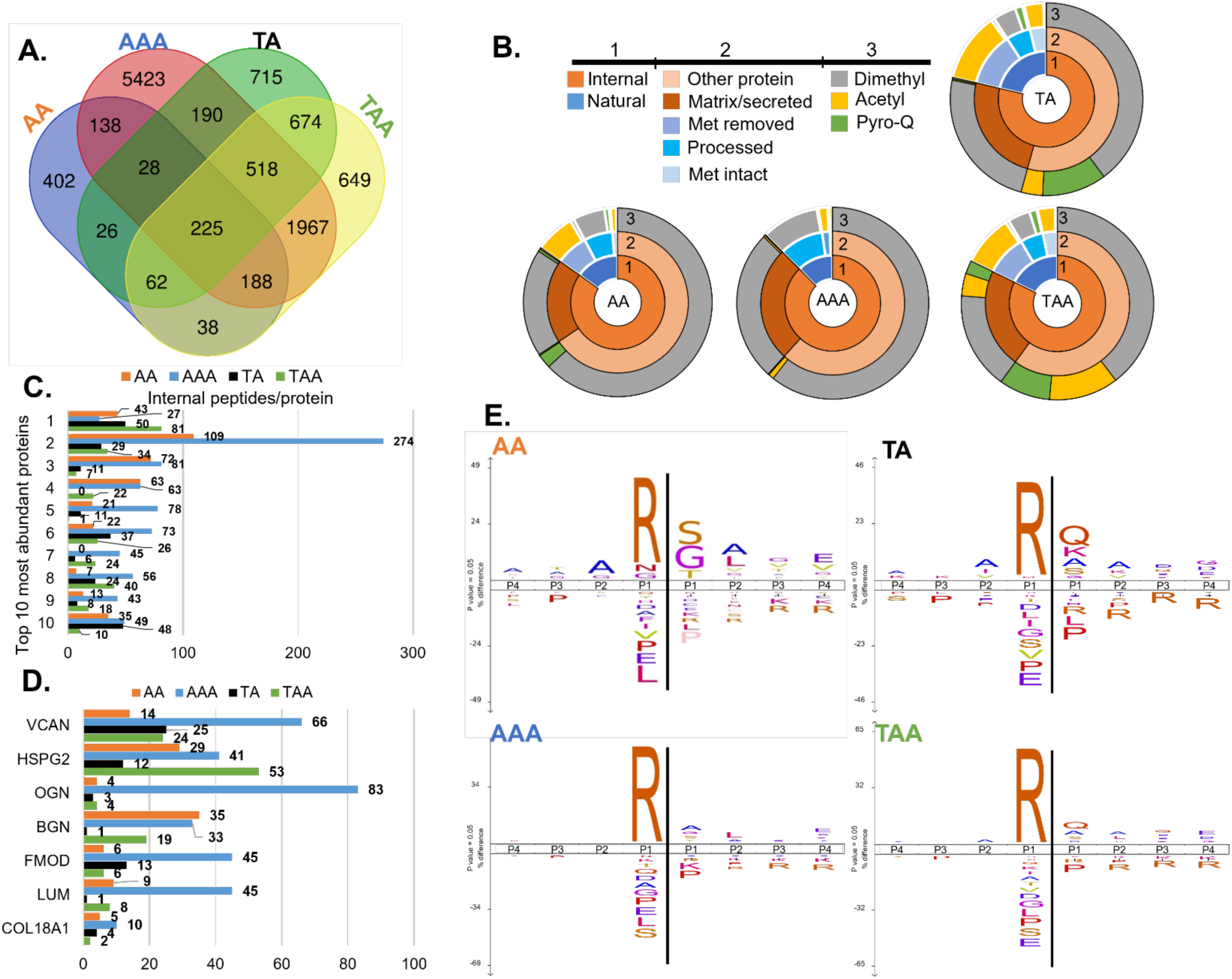
Dimethyl TAILS reveals increased proteoglycan proteolysis in aortas. A. Venn diagram of the internal peptides identified in each tissue cohort showing shared and unique cleavages. B. Sunburst plots from blocked peptides from each of the tissue cohorts shows broadly similar N-terminomes. C. The number of cleavage sites/ protein for the top 10 most abundant proteins in each aortic cohort (Gene symbols for each are in Figure S3D). D. The number of cleavage sites/ proteoglycan identified in in each aortic cohort. E. IceLogo plots from each group indicating the frequently observed amino acids around the cleavage sites (vertical line); letter size reflects the relative frequency with which the amino acid was observed.

The dimethyl degradome included peptides from 7 proteoglycans, each with varying numbers of specific cleavages (Figure 3D) and AAAs or TAAs contained the most cleavage sites in each proteoglycan except biglycan. AAAs had the most cleavage sites in versican (66), mimecan (83), fibromodulin (45), lumican (45), and collagen α1 (XVIII) (10) (Figure 3D), whereas TAAs had the most cleavage sites in perlecan (HSPG2) and AAs had the most cleavage sites in BGN. More cleavage sites in all proteoglycans were identified in AAAs than in donor aortas.

IceLogo plots, which summarize the frequency of occurrence of amino acids around the identified cleavage sites in relation to their natural abundance indicated an overall Arg-C-like preference, i.e., with arginine preferred at the P1 residue (the C-terminal residue resulting from scissile bond cleavage (Schechter and Berger nomenclature[20])) in all cohorts (Figure 3E). When iceLogo plots were re-constructed using all peptides without arginine at P1, AAAs showed a preference for valine, isoleucine, and phenylalanine at P1, whereas TAs showed a preference for glycine at P1 (Figure S2); such IceLogo plots for TAAs and TAs differed by presence of an acidic residue (aspartic acid (D) or glutamic acid (E)) in the P2’ position in TAAs (Figure S2).

### Quantitative analysis of proteolytically cleaved peptides between human aorta cohorts

To determine whether the cohort proteomes had distinct compositional profiles, all proteins identified in each cohort by at least two peptides were first used for principal component analysis (PCA). PCA accounted for 70% of the data (component 1=48.5%, component 2=25%), showing excellent cohort segregation (Figure S3A). Unbiased hierarchical clustering using all quantifiable proteins showed that TAAs resembled each other more than AAAs and vice versa (Figure S3B). Of 1629 proteins identified in TAAs and AAAs, 649 (39.8%) overlapped between the two cohorts (Figure S3C). Compared to control aortas (combined AAs and TAs), 358 and 290 proteins were exclusively identified in AAAs and TAAs, respectively. Each cohort showed substantial contribution from the cytosolic proteins actin and vimentin, circulating proteins, and matrix proteins, such as collagens (collagen I and collagen VI), which comprised over 60% of the total ion intensity in TAAs and AAAs (Figure S3D). However, comparing protein class abundance from AAAs with TAAs, blood-related proteins and cytosolic proteins had the greatest respective contributions (Figure S3E). Pathway enrichment analysis via PantherDB.org revealed blood coagulation as a major contributor in AAAs whereas TAAs had greater contributions from cytokine and chemokine mediated inflammation pathways, B-cell activation, and glycolysis (Figure S3F).

To determine changes in abundance of proteolytically cleaved peptides, we undertook quantitative degradomic analysis on four paired cohorts, i) AAA vs TAA ii) AAA vs AA iii) TAA vs TA and iv) AA vs TA using 8-plex iTRAQ TAILS (Fig. 1A). N-terminomes from iTRAQ labeled cohorts were classified by modification as in the dimethyl labeling strategy (Figure 2B) but only N-terminal iTRAQ labeled internal peptides were included in the degradome since naturally modified peptides cannot be assigned by individual sample in multiplexed experiments. PCA plots and unsupervised hierarchical clustering showed consistent sample segregation by cohort (Figures 4A, B, S4, S5, and S6) other than one TA sample which clustered with the TAAs (Figure S6A). We further annotated the N-terminally blocked peptides as natural and internal (Figures 4C, S4, S5, S6). Cellular component analysis of internal peptides showed that ECM proteins were cleaved as frequently as cellular proteins in all cohorts (Figures 4D, S5), together accounting for 80% of the proteolytically cleaved peptides. Statistical analysis of cohort pairs (Figures 4E, S4, S5, 6D) found most peptides with significant abundance changes in AAAs vs TAAs (Figure 4E). No statistically significant peptides were identified in TAA vs TA comparison, likely owing to the single TA outlier (Figure S6D).

**Figure 4.**
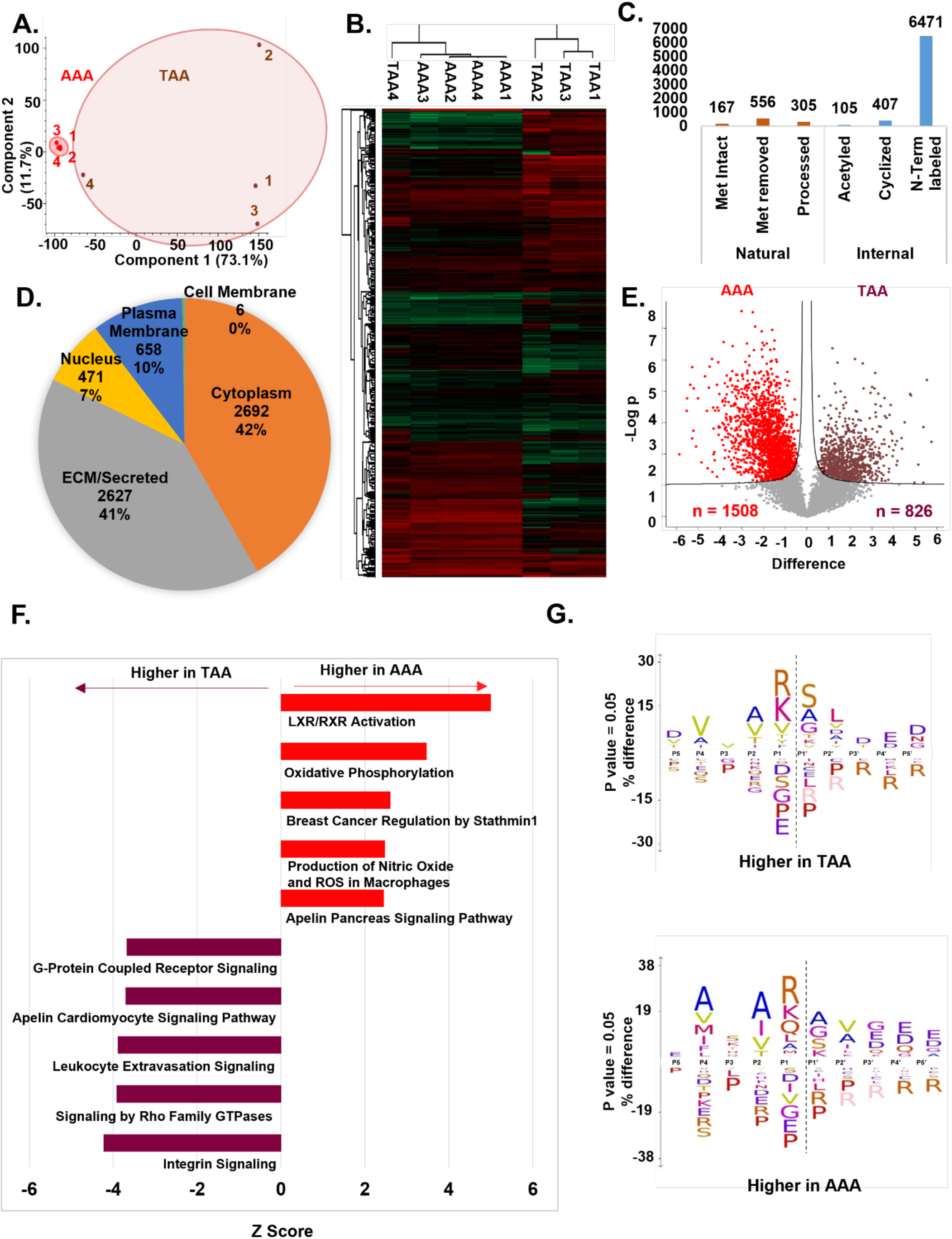
Contrasting degradomes of thoracic and abdominal aortic aneurysms: A. Principal Component analysis based on internal peptides of 8-plex iTRAQ-TAILS of TAAs and AAAs showed that the first and second components segregate the cohorts and account for 50% and 23.5% of the data, respectively. B. Heat map of significant peptide abundances (ANOVA, FDR < 0.05) after unsupervised hierarchical clustering shows clustering by cohort other than TAA4, which appears to be an outlier. C. Peptide count annotations of the N-terminome by modification and position of the identified peptides mapped on the mature protein after 8-plex iTRAQ-TAILS D. Pie chart showing the cellular component distribution of the proteolytically cleaved peptides in TAAs and AAAs from iTRAQ-TAILS. E. Volcano plot of the internal peptides from TAAs and AAAs identifying quantitative differences between their substrate degradomes. F. Top 5 canonical pathways associated with each cohort (based on Z score) as generated by Ingenuity Pathway Analysis (IPA) providing an overview of the major biological differences between TAAs and AAAs. G. IceLogo plot of statistically significant peptides in TAA and AAA substrate degradomes obtained by comparing the amino acid frequency around the cleavage sites (vertical dotted line) against frequency in the human proteome database.

Ingenuity pathway analysis (IPA) of protein abundance changes identified in preTAILS analysis of each cohort pair comparison suggested activation of the LXR/RXR pathways in AAA when compared to TAA and AA cohorts (Figures 4F, S5F), but also showed higher activation of this pathway in the AA cohort compared to the TA cohort (Figure S4F). Integrin signaling showed the highest predicted activation levels in TAAs compared to AAAs and in TAs compared to AAs (Figures 4F, S4F), suggesting fundamental cellular differences between abdominal and thoracic aortas. IceLogo plots using the significant proteolytic peptides from each cohort showed only minor differences with the highest preference for arginine at the P1 position in all groups (Figures 4F, S4G and S5G). The only clear difference between cohorts was the secondary preference for aspartic acid in the P1 position in the TA cohort compared to the AA cohort (Figure S4G).

### The aortic wall protease degradome

To define protease degradomes we utilized shotgun-like analysis of the dimethyl dataset (pre-TAILS samples) and included proteases identified in at least two replicates within a cohort. No proteases were detected in iTRAQ-TAILS data likely because ion stacking after multiplexing masks peptides from low abundance proteins such as proteases. Among the 90 proteases identified, matrix metalloproteases (MMPs) (n=27) and serine proteases (n=38) were most numerous (Figure 5A). 32 proteases were identified only in AAAs or TAAs but not in TAs or AAs. No MMPs were found in AAs, and 12 serine proteases were detected exclusively in either the TAAs or AAAs (Figure 5A). 9 proteases (DPP9, TPSD1, PRSS23, PRSS1, PSMA2, PSMA8, PSMB2, NPEPL1, and NPEPPSL) were seen only in non-diseased aortas and 5 of the 9 were found in at least 3/4 TAs (Figure 5A).

**Figure 5.**
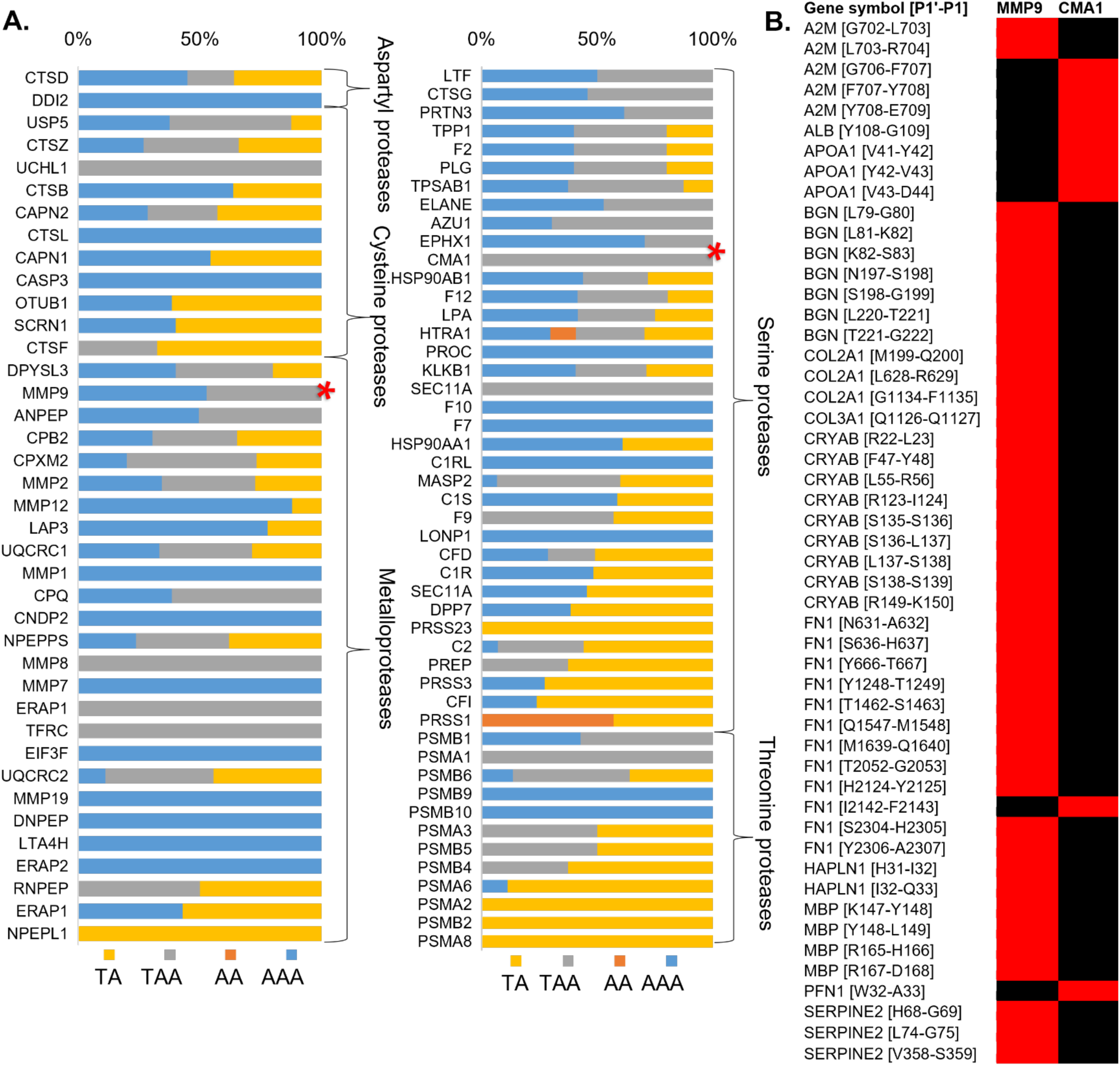
Protease degradomes of normal and aneurysmal aorta. A. Proteases identified in aortic tissue and the frequency of their detection in each cohort. Asterisks indicate CMA1 and MMP9, the two proteases chosen for reverse degradomics. B. CMA1 or MMP9 cleavage sites (red boxes in each column) identified in the aortic substrate degradome matching those previously reported (in MEROPS and TopFIND 4.0) illustrate distinct sites of cleavage.

We utilized TopFINDer 4.0 [21] to attribute internal peptides to the activity of protease(s) previously shown to cleave at a particular site. 1,042 cleavages in the aortic substrate degradome were assigned to 83 proteases by TopFINDer (Figure S7A, B). Of these proteases, 28 were experimentally identified in the aortic protease degradome (Figures 5A, S7B) and together accounted for 376/1,042 cleavage sites matched to proteases in TopFINDer. Specific metalloproteases (MMP 1, 7, 8, and 9), serine proteases (CMA1, CTSG, F10, and C1RL) and cysteine proteases (CTSL and CASP3) were identified only in TAAs and AAAs.

MMP9 was detected in 12/16 AAAs and 4/5 TAAs whereas CMA1 was found exclusively in all 5 TAAs. 42 previously reported MMP9 cleavage sites and 9 CMA1 cleavage sites matched peptides found in the aorta degradomes (Figure 5B). Known substrates of these proteases that we identified included biglycan and several collagens (cleaved by MMP9), α2-macroglobulin (a broad-spectrum protease substrate and inhibitor) and FN1, which was cleaved by both MMP9 and CMA1 (Figure 5B), as well as distinct substrates of each. This paucity of knowledge about aorta-specific substrates, prompted us to investigate MMP9 and CMA1 in a reverse degradomics analysis, where we digested aorta tissue with each protease, followed by TAILS (Figure 1A), with the goal of detecting such experimentally determined cleavages in aorta substrate degradomes.

### Aorta-specific proteolytic profile of CMA1

Aortic digestion with CMA1 followed by duplex dimethyl-TAILS identified 164 unique internal peptides with significantly higher abundance over control digests of which 90 peptides from 36 ECM/secreted proteins were considered further since CMA1 is a secreted protease (Figure 6A). IceLogo analysis of the 90 sites showed broadly similar preferences with prevalence of Phe, Tyr, Arg, and Leu at the P1 position and prevalence of Ser, Ala, and Asp at P2′ (Figure 6A). Previously, digestion of a 9-residue peptide library had demonstrated chymotrypsin-like specificity of CMA1 with preference for Phe, Tyr, Trp, and Leu at the P1 position and acidic residues at the P2′ position [22]. 11/90 cleavage sites matched protease cleavages in the TopFIND 4.0 search, which were attributed to MMP2, MMP9, MMP12, MMP13, and ADAMTS4 and ADAMTS5 (Supplemental data file 2), suggesting overlapping substrate and site specificity or potential activation of the MMPs by CMA1.

**Figure 6.**
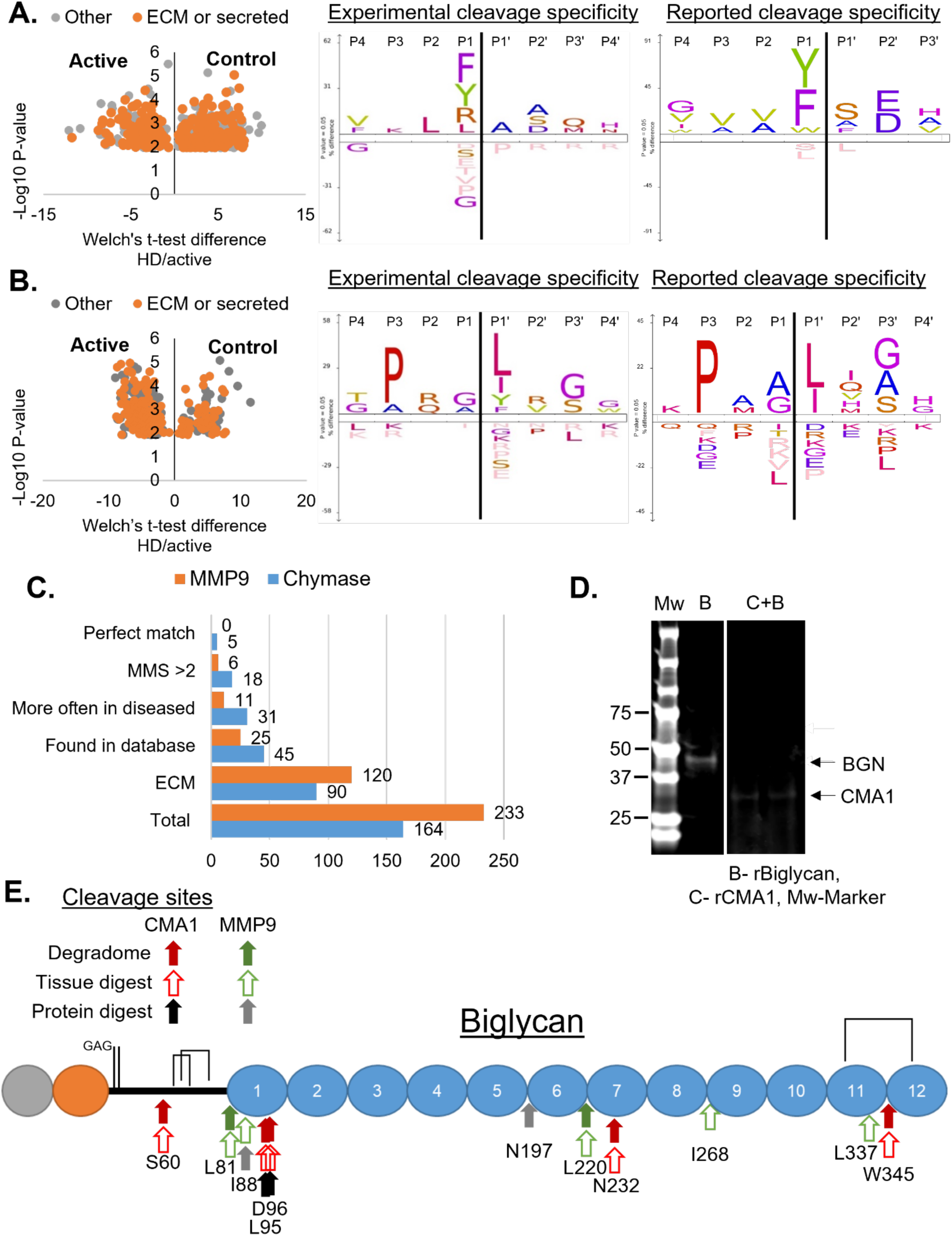
ATOMS analysis for identification of CMA1 and MMP9 substrates and their cleavage sites finds novel aortic aneurysm relevant substrates. A,B. (left) Volcano plot of significant peptides in CMA1 (A) or MMP9 (B) digests. (Right) IceLogo plots for CMA1 (A) and MMP9 (B) cleavages in ECM and secreted substrates were compared to their reported cleavage site specificity. C. Cleavages were heirarchically sorted based on substrate localization (ECM), presence in the aortic substrate degradome, detection in a disease cohort, motif matching score (MMS) greater than 2 (MMS>2) and if it matches the predicted motif perfectly (Perfect match) D. Biglycan cleavage by CMA1. Coomassie blue stained gel after reducing SDS-PAGE showing less recombinant biglycan in the presence of CMA1 (arrows indicate the molecular mass of each). Molecular weight markers are shown on the left. E. Domain structure of biglycan showing the CMA1 and MMP9 cleavages identified by ATOMS, and the overlap with sites identified in aortic protein digest with each protease and the aortic substrate degradome.

The 90 CMA1 ECM cleavage sites matched 45 cleavages in the aorta degradome (Figure 6C, Supplemental data file 2) and over half (31/45, 69 %) occurred more often in AAAs or TAAs than in non-diseased tissue. These sites were compared to the known CMA1 protease specificity by a motif matching scoring system, allotting 2 points for Phe, Tyr or Trp at P1,1 for Leu at P1 and 1 for acidic residues at P2′ (Figure 6A). 18/31 sites had a motif matching score >2 and 5/18 cleavage sites from 5 ECM proteins (proteoglycans VCAN, OGN, PRELP, BGN, and COL6A3) matched the chymotrypsin-like preference of CMA1 at both P1 and P2’ positions (Figure 6C, Table 1). PRELP, VCAN, and BGN are additionally cleaved within 1 amino acid of the observed CMA1 sites by other proteases including ADAMTS4, ADAMTS5 and MMP13, suggesting that these CMA1 cleavages occur within proteolytically susceptible regions.

**Table 1.** CMA1 and MMP9 cleavage sites with statistically significant abundance in aneurysms. *Both= found in both iTRAQ and dimethyl TAILS, *= only found in the active protease digest*

|  |  |  |  |  | Dimethyl |  |  |  | iTRAQ |  |  |  |  |
| --- | --- | --- | --- | --- | --- | --- | --- | --- | --- | --- | --- | --- | --- |
| Protease | Peptide annotation | N-terminal labeling method | Motif Matching Score | Ratio of peptide abundance from active/inactive | AA | AAA | TA | TAA | AA vs TA | AAA vs AA | AAA vs TAA | TAA vs TA | Other proteases cleaving this site |
| CMA1 | COL6A3 [F1951-T1952] | Dimethyl | 3 | 81.46 | 0% | 84% | 0% | 40% |  |  |  |  |  |
| CMA1 | PRELP [F339-H340] | Both | 3 | 7.17 | 0% | 78% | 63% | 70% | -0.29 |  |  | -0.20 | MMP13;ADAMTS4/5 |
| CMA1 | OGN [Y219-L220] | Both | 3 | 12.21 | 0% | 13% | 13% | 70% | -0.07 |  |  |  | ADAMTS4 |
| CMA1 | VCAN [Y189-E190] | Dimethyl | 3 | 13.63 | 0% | 0% | 63% | 100% |  |  |  |  |  |
| CMA1 | BGN [Y344-W345] | Both | 3 | 100* | 0% | 3% | 50% | 50% | 0.05 |  |  |  | MMP13 |
| MMP9 | AEBP1 [R732-Y733] | Dimethyl | 3 | 100* | 0% | 97% | 0% | 60% |  |  |  |  |  |
| MMP9 | AEBP1 [R732-Y733] | Dimethyl | 3 | 100* | 0% | 97% | 0% | 60% |  |  |  |  |  |
| MMP9 | VTN [N242-I243] | Both | 3 | 100* | 0% | 3% | 0% | 0% | 0.88 |  |  |  |  |
| MMP9 | COL6A2 [S287-L288] | Dimethyl | 3 | 100* | 0% | 3% | 0% | 0% |  |  |  |  |  |
| MMP9 | HTRA1 [E210-L211] | Dimethyl | 3 | 100* | 0% | 3% | 0% | 0% |  |  |  |  |  |
| MMP9 | VTN [L195-I196] | Dimethyl | 3 | 100* | 0% | 3% | 0% | 0% |  |  |  |  |  |

### Experimentally defined aorta-specific proteolytic profile of MMP9

Duplex dimethyl-TAILS identified 233 internal N-terminally labeled peptides with higher abundance in MMP9 digest of aortic proteins. Peptides from ECM/secreted proteins comprised 52% (120/233) of the peptides and were considered further, since MMP9 is a secreted protease (Figure 6B, C). IceLogo analysis of these sites demonstrated dominance of Pro at the P3 position, with Leu and Ile preferred at P1ʹ, consistent with reported MMP9 preferences [23]. 6 sites matched known MMP9 cleavage sites in fibronectin, collagen α1 (III) and biglycan (Supplemental data file 2). 25 of the 120 ECM peptides were identified in the aortic substrate degradome, although not the 6 previously reported MMP9 cleavages. Cleavage site at L^155^-Y^156^ was previously attributed to MMP12, MMP3 and ADAMTS4, and at N^222^-L^223^ to MMP8.

11 MMP9 cleavages had higher abundance in aneurysms. Analysis using the motif-matching scoring system allotting 2 points for Pro at P3, 1 point for Leu/ Ile at P1 and 1 point for Ser/Gly at P3′ (Figure 6C), identified 6/11 sites with a motif matching score of >2, (Figure 6C, Table 1). None of the six sites were previously reported (Table 1) [24–26]. Two of the six sites were identical but arose from peptides with different C-terminal ends. This cleavage site in adipocyte enhancer–binding protein 1 was found in over half (6/10) of the TAA dimethyl samples and all but one (35/36) of the AAAs. The remaining 5 cleavage sites occurred in only the AAAs and were from vitronectin, collagen α2(VI), and serine protease HtrA1. The vitronectin peptide indicating cleavage at N^242^-I^243^ was identified in the iTRAQ comparison of TA *vs* AA but was not statistically significant.

### Orthogonal validation of biglycan as a direct CMA 1 and MMP9 substrate

Since MMP9 and CMA1 may modify activity of other proteases, we used <u>A</u>mino-<u>T</u>erminal <u>O</u>riented <u>M</u>ass spectrometry of <u>S</u>ubstrates (ATOMS) [27], to define CMA1 and MMP9 cleavage sites in biglycan using recombinant proteins. Biglycan cleavage sites were identified in both MMP9 and CMA1 aorta digests and in the aortic substrate degradome. Biglycan is a known MMP9 substrate and matching cleavage sites were identified in the aortic degradomes [28, 29] (Figure 5B). CMA1 was not previously known to cleave biglycan, despite a preference for proteoglycans [22, 30]. Digestion of recombinant biglycan with recombinant CMA1 or MMP9 was was validated by western blot of the digests (Figure 6D). ATOMS identified two CMA-1 cleaved biglycan peptides indicating cleavage at L^95^-D^96^ and D^96^-L^97^ which were absent in tissue and tissue digest degradomes (Figure 6E, Table 2).

**Table 2.** Biglycan peptides of interest. NF= Not found, Both= found in the iTRAQ and dimethyl database, += found with N-terminal and C-terminal peptides, *= only found in the protease active

|  |  |  |  |  | Dimethyl |  |  |  | iTRAQ |  |  |  |  |
| --- | --- | --- | --- | --- | --- | --- | --- | --- | --- | --- | --- | --- | --- |
| Protease | Gene [P1-P1']<br>Cleavage site | BGN digest ratio<br>(active/inactive) | Tissue digest ratio<br>(active/inactive) | Found in<br>aortic<br>substrate<br>degradomes | AA | AAA | TA | TAA | AAv<br>TA | AAAv<br>AA | AAAv<br>TAA | TAAv<br>TA | Known<br>proteases |
| MMP9 | BGN [196-197] | 100 | NF | NF |  |  |  |  |  |  |  |  |  |
| MMP9 | BGN [87-88] | 100 | 7.271 | NF |  |  |  |  |  |  |  |  |  |
| MMP9 | BGN [219-220] |  | 6.321 | iTRAQ |  |  |  |  |  | 1.10 |  |  | MMP3/9/12/13 |
| MMP9 | BGN [336-337] |  | 6.163 | NF |  |  |  |  |  |  |  |  |  |
| MMP9 | BGN [80-81] |  | 5.038 | NF |  |  |  |  |  |  |  |  | MMP3/9/13 |
| MMP9 | BGN [267-268] |  | 4.132 | NF |  |  |  |  |  |  |  |  |  |
| CMA1 | BGN [95-96] | 100 | 4.998 | Both | 0% | 81% | 13% | 90% |  |  |  | Found |  |
| CMA1 | BGN [94-95] | 100 | 5.43 | Both | 0% | 69% | 0% | 50% | 1.72 | 2.42 |  | 0.64 |  |
| CMA1 | BGN<br>[Y344-W345] |  | 100* | Both+ | 0% | 3% | 50% | 50% | 0.05 |  |  |  | MMP13 |
| CMA1 | BGN [231-232] |  | 44.435 | Both | 0% | 63% | 0% | 0% | 1.37 |  |  |  | MMP12 |
| CMA1 | BGN [Y59-S60] |  | 2.255 | Both | Found |  |  |  |  | 0.445 |  |  |  |

MMP9 digestion of biglycan identified two peptides indicating cleavage sites at I^88^-S^89^ and N^197^-S^198^, the latter previously known (Table 2). Of 6 MMP9 cleavage sites in biglycan, the L^220^-T^221^ site was found in both in the aorta degradome and the aortic digest but not in the recombinant biglycan digest, and the I^88^-S^89^ site was identified in both the tissue digest and the recombinant biglycan digest (Figure 6E, Table 2). Notably, most observed cleavages with each protease were located between leucine-rich repeats. These examples demonstrate how forward and reverse degradomics can be integrated to detect novel activities, provide disease context to those previously known and clarify the activity of specific proteases in aortic disease. For user-friendly access to proteolytic data from forward and reverse degradomics using either protein name or UniProt ID we built the web interface named Database of Identified Cleavage Sites Endemic to Disease States (DICED, website) (Figure S8).

## Discussion

Here we demonstrate that integrated forward and reverse degradomics provides previously unavailable insights on proteolysis in healthy and aneurysmal aortas and emerges as a feasible strategy for investigation of proteolytic mechanisms in cardiovascular disease. The aortic degradomes represent an important step forward for aneurysm research because they provide a tissue and disease-specific resource. In addition to compositional and biological process differences in the control and aneurysm aorta, the work has shown that proteoglycans and other ECM components undergo disproportionately greater proteolysis in aneurysms, thus having potentially greater influence on aneurysms than previously suspected.

Prior to invention of terminomics methods such as TAILS, proteolytic events were typically identified by biochemical analysis and extrapolated to a tissue context. In the current approach, proteolytic events are first detected within the aortic tissue context for unequivocal evidence of cleavage occurring locally, and constituting an ensemble of all proteolytic events, i.e., the degradome. The proteolytic landscape thus defined facilitates biomarker prospecting by providing new peptides which have a functional overlay of proteolysis. For example, neo-epitope antibodies specifically recognizing aggrecan and versican neo-N- or C-termini resulting from proteolysis were used in prior analysis of vascular disease, including aortic aneurysms [31–36]. Another important utility of the aortic degradome is to define which cleavages result from specific proteases such as MMP9 and CMA1 to better target them in disease. The approach can be used to define the role of any protease.

Proteases rarely have a stringent sequence preference which can be used for cleavage site prediction and many have additional determinants of activity such as exosites that bind the substrate distant from the cleaved peptide bond [37], substrate post-translational modifications such as phosphorylation and glycosylation [38], mechanical unfolding of the substrate [39] or allostery. Similarly, substrate site preferences, elucidated for many proteases using peptide libraries, cannot accurately predict novel sites either.

Although proteomics has made enormous contributions to understanding vascular disease, proteolytic activity was previously investigated indirectly or by testing for candidate cleavages using neo-epitope antibodies that each detect only a single cleavage site [33, 36, 40–42]. For example, prior studies which specifically addressed proteolysis in aortic aneurysms identified proteins released by digestion with ADAMTS5 or MMP12 [33, 34]. Substrates of MMP3, MMP14 and MMP12 in radial artery digests were sought by analysis of released proteins and semi-tryptic peptides [29]. In such analysis, identified proteins could be released from complexes without undergoing proteolysis. Without N-terminal labeling and by relying on semi-tryptic sites, trypsin-like cleavages (with P1 Arg residue), which as we show, constitute the majority, would be missed.

Only a handful of TAA and AAA biomarkers have been identified to date [43, 44]. Circulating markers such as D-dimer [45] and C-reactive protein [46] may originate outside the aortic wall and are not specific for aortic aneurysms. ECM proteins whose levels are altered in aneurysms such as collagens XI and V [47] are not yet translated into serum biomarkers, although a promising association was noted between fibrillin fragments and TAA dissection [48]. Dimethyl TAILS identified numerous proteases which were only found in TAAs and not the TAs cohort. Proteases could themselves be used as biomarkers however, their proteolytic products would indicate both the presence of the protease and its activity. The iTRAQ degradome identified over 400 internal peptides with significant changes between the AAAs and controls. These could also be used as biomarkers although most of the inferred cleavage sites from these peptides are not currently annotated to a specific protease. Additionally, the dimethyl analysis identified 239 previously unannotated internal peptides which were found only in the diseased tissues and not in the controls. Thus, the aortic degradome offers a useful resource for potential new biomarkers.

Mutations in the novel MMP9 substrate AEBP1 associate with Ehlers-Danlos syndrome and development of AAA [49]. AEBP1 was previously suggested as a potential target of MMP9 although direct cleavage was not shown [34]. The R^732^-Y^733^ cleavage site in AEBP1 was identified in 97% of the AAA samples using dimethyl analysis, 60% of the TAA samples and none of the control samples. Furthermore, we identified 5 novel CMA1 substrates and cleavage sites which occurred more often in the aneurysms. Although proteoglycans have long been suspected as substrates of CMA1 [22], specific cleavage sites had yet to be identified. Two biglycan cleavages at L^94^-D^95^ and D^95^-L^96^ in the aortic degradome were also identified after tissue and biglycan digestion with CMA1. Peptides indicating these cleavage sites were present in 81% of AAAs and 90% of TAAs, and 69% of the AAAs and 50% of the TAA,s respectively, yet only one of the control aortas. As AEBP1 and BGN are ECM proteins, there is high possibility that their cleavage products could translocate to the circulation to provide disease biomarkers detectable by targeted LC-MS/MS MS-based assays or neoepitope antibodies. It is notable, however, that the experimentally obtained CMA1 and MMP9 cleavages account for only a fraction of the degradome, implying that numerous other proteases participate in aortic wall remodeling.

## NOVELTY AND SIGNIFICANCE

### What is known?

- The aortic wall is suspected to undergo extensive remodeling in aneurysms, but there is little current experimental evidence for this.
- Proteases that degrade extracellular matrix are causally implicated in aortic aneurysms and some, such as MMP9, are considered to be drug targets [50, 51].

### What new information does this article contribute?

- The present work contributes a database of thousands of cleavage sites in numerous proteins in the normal aorta and aortic aneurysms, providing proteome-wide molecular definition of vascular remodeling.
- Specific differences in proteolytic landscapes were detected in TAAs and AAAs, and potential biomarkers were uncovered.
- Targets of two of the most consistently identified proteases, MMP9 and CMA1, were identified in the aortic wall.
- Integrated forward-reverse degradomics can elucidate the activity profile of any protease in cardiovascular disease.

### Summary of Novelty and Significance

Although proteases are frequently associated with TAAs/AAAs by mRNA detection or immunostaining, neither is a proper surrogate for their activity, since many are synthesized as zymogens that require activation and have ambient inhibitors. The only true measure of ECM proteolysis is a cleaved peptide bond, but a complete landscape of aortic proteolysis, i.e., molecules cleaved and cleavage sites therein (substrate degradome), and proteases generating the cleavages (protease degradome) remained largely undefined. By precisely identifying protein cleavage sites in an unbiased manner in non-diseased and aneurysmal aortic tissue, we demonstrate that the proteolytic landscape of human aortic aneurysms is vast and shows distinct differences between ascending thoracic and abdominal aortic aneurysms. Specific substrates and cleavage sites of MMP9 and CMA1, proteases which were consistently detected in aortic wall, defined their role in vascular wall breakdown mechanistically. The work establishes a new paradigm for fully defining proteolysis at a molecular level on a proteome-wide scale and for elucidating the specific roles of individual proteases not only in aneurysms, but any cardiovascular disorder, to provide disease biomarkers and new targetable mechanisms.

## Acknowledgements

This work is dedicated to the late Professor Ulrich auf dem Keller. We are indebted to him and Professor Christopher M. Overall for encouragement, guidance and key resources in degradomics.

## Nonstandard abbreviations and nonstandard acronyms

AA: Abdominal aorta
AAA: Abdominal aortic aneurysm
ATOMS: Amino-Terminal Oriented Mass Spectrometry of Substrates
ECM: Extracellular matrix
iTRAQ: Isobaric tags for relative and absolute quantitation
LC-MS/MS: Liquid chromatography tandem mass spectroscopy
MMP: Matrix metalloproteinase
SMC: Smooth muscle cell
TA: Thoracic aorta
TAA: Thoracic aortic aneurysm
TAILS: Terminal amine isotopic labeling of substrates

## References

1. Humphrey, J.D., et al., Cell biology. Dysfunctional mechanosensing in aneurysms. Science, 2014. 344(6183): p. 477-9.

2. Halushka, M.K., et al., Consensus statement on surgical pathology of the aorta from the Society for Cardiovascular Pathology and the Association For European Cardiovascular Pathology: II. Noninflammatory degenerative diseases - nomenclature and diagnostic criteria. Cardiovasc Pathol, 2016. 25(3): p. 247–57.

3. Michel, J.B., et al., Novel aspects of the pathogenesis of aneurysms of the abdominal aorta in humans. Cardiovasc Res, 2011. 90(1): p. 18–27.

4. Osborne, J.D., et al., Annotating the human genome with Disease Ontology. BMC Genomics, 2009. 10 **Suppl 1**: p. S6.

5. Hendel, A., L.S. Ang, and D.J. Granville, Inflammaging and proteases in abdominal aortic aneurysm. Curr Vasc Pharmacol, 2015. 13(1): p. 95–110.

6. Liu, B., et al., Pathogenic mechanisms and the potential of drug therapies for aortic aneurysm. Am J Physiol Heart Circ Physiol, 2020. 318(3): p. H652–h670.

7. Mougin, Z., et al., ADAMTS Proteins and Vascular Remodeling in Aortic Aneurysms. Biomolecules, 2021. 12(1).

8. Rabkin, S.W., The Role Matrix Metalloproteinases in the Production of Aortic Aneurysm. Prog Mol Biol Transl Sci, 2017. 147: p. 239–265.

9. Lopez-Otin, C. and C.M. Overall, Protease degradomics: a new challenge for proteomics. Nat Rev Mol Cell Biol, 2002. 3(7): p. 509–19.

10. auf dem Keller, U., et al., Systems-level analysis of proteolytic events in increased vascular permeability and complement activation in skin inflammation. Sci Signal, 2013. 6(258): p. rs2.

11. Bhutada, S., et al., Forward and reverse degradomics defines the proteolytic landscape of human knee osteoarthritic cartilage and the role of the serine protease HtrA1. Osteoarthritis Cartilage, 2022.

12. Prudova, A., et al., TAILS N-terminomics of human platelets reveals pervasive metalloproteinase-dependent proteolytic processing in storage. Blood, 2014. 124(26): p. e49–60.

13. Hattori, N., et al., Pericellular versican regulates the fibroblast-myofibroblast transition: a role for ADAMTS5 protease-mediated proteolysis. J Biol Chem, 2011. 286(39): p. 34298–310.

14. Mead, T.J., et al., ADAMTS9-Regulated Pericellular Matrix Dynamics Governs Focal Adhesion-Dependent Smooth Muscle Differentiation. Cell Rep, 2018. 23(2): p. 485–498.

15. Sakalihasan, N., R. Limet, and O.D. Defawe, Abdominal aortic aneurysm. Lancet, 2005. 365(9470): p. 1577-89.

16. Tamarina, N.A., et al., Expression of matrix metalloproteinases and their inhibitors in aneurysms and normal aorta. Surgery, 1997. 122(2): p. 264–71; discussion 271-2.

17. Li, T., et al., Serum matrix metalloproteinase-9 is a valuable biomarker for identification of abdominal and thoracic aortic aneurysm: a case-control study. BMC Cardiovasc Disord, 2018. 18(1): p. 202.

18. Sun, J., et al., Critical role of mast cell chymase in mouse abdominal aortic aneurysm formation. Circulation, 2009. 120(11): p. 973–82.

19. Swedenborg, J., M.I. Mäyränpää, and P.T. Kovanen, Mast cells: important players in the orchestrated pathogenesis of abdominal aortic aneurysms. Arterioscler Thromb Vasc Biol, 2011. 31(4): p. 734–40.

20. Schechter, I. and A. Berger, On the size of the active site in proteases. I. Papain. Biochem Biophys Res Commun, 1967. 27(2): p. 157–62.

21. Fortelny, N., et al., Proteome TopFIND 3.0 with TopFINDer and PathFINDer: database and analysis tools for the association of protein termini to pre- and post-translational events. Nucleic Acids Res, 2015. 43(Database issue): p. D290-7.

22. Andersson, M.K., et al., The extended substrate specificity of the human mast cell chymase reveals a serine protease with well-defined substrate recognition profile. Int Immunol, 2009. 21(1): p. 95–104.

23. Eckhard, U., et al., Active site specificity profiling of the matrix metalloproteinase family: Proteomic identification of 4300 cleavage sites by nine MMPs explored with structural and synthetic peptide cleavage analyses. Matrix Biol, 2016. 49: p. 37–60.

24. Rawlings, N.D. and A.J. Barrett, MEROPS: the peptidase database. Nucleic Acids Res, 1999. 27(1): p. 325–31.

25. Rawlings, N.D., et al., The MEROPS database of proteolytic enzymes, their substrates and inhibitors in 2017 and a comparison with peptidases in the PANTHER database. Nucleic Acids Res, 2018. 46(D1): p. D624–d632.

26. Rawlings, N.D. and A. Bateman, How to use the MEROPS database and website to help understand peptidase specificity. Protein Sci, 2021. 30(1): p. 83–92.

27. Doucet, A. and C.M. Overall, Amino-Terminal Oriented Mass Spectrometry of Substrates (ATOMS) N-terminal sequencing of proteins and proteolytic cleavage sites by quantitative mass spectrometry. Methods Enzymol, 2011. 501: p. 275–93.

28. Genovese, F., et al., Biglycan fragmentation in pathologies associated with extracellular matrix remodeling by matrix metalloproteinases. Fibrogenesis Tissue Repair, 2013. 6(1): p. 9.

29. Stegemann, C., et al., Proteomic identification of matrix metalloproteinase substrates in the human vasculature. Circ Cardiovasc Genet, 2013. 6(1): p. 106–17.

30. Thorpe, M., et al., Extended cleavage specificities of mast cell proteases 1 and 2 from golden hamster: Classical chymase and an elastolytic protease comparable to rat and mouse MCP-5. PLoS One, 2018. 13(12): p. e0207826.

31. Barallobre-Barreiro, J., et al., Extracellular Matrix in Heart Failure: Role of ADAMTS5 in Proteoglycan Remodeling. Circulation, 2021. 144(25): p. 2021–2034.

32. Cikach, F.S., et al., Massive aggrecan and versican accumulation in thoracic aortic aneurysm and dissection. JCI Insight, 2018. 3(5): p. e97167.

33. Didangelos, A., et al., Novel role of ADAMTS-5 protein in proteoglycan turnover and lipoprotein retention in atherosclerosis. J Biol Chem, 2012. 287(23): p. 19341–5.

34. Didangelos, A., et al., Extracellular matrix composition and remodeling in human abdominal aortic aneurysms: a proteomics approach. Mol Cell Proteomics, 2011. 10(8): p. M111 008128.

35. Fava, M., et al., Role of ADAMTS (A Disintegrin and Metalloproteinase With Thrombospondin Motifs)-5 in Aortic Dilatation and Extracellular Matrix Remodeling. Arterioscler Thromb Vasc Biol, 2018.

36. Suna, G., et al., Extracellular Matrix Proteomics Reveals Interplay of Aggrecan and Aggrecanases in Vascular Remodeling of Stented Coronary Arteries. Circulation, 2018. 137(2): p. 166–183.

37. Foulcer, S.J., et al., Determinants of versican-V1 proteoglycan processing by the metalloproteinase ADAMTS5. J Biol Chem, 2014. 289(40): p. 27859–73.

38. Tagliabracci, V.S., et al., Dynamic regulation of FGF23 by Fam20C phosphorylation, GalNAc-T3 glycosylation, and furin proteolysis. Proc Natl Acad Sci U S A, 2014. 111(15): p. 5520-5.

39. Wu, T., et al., Force-induced cleavage of single VWFA1A2A3 tridomains by ADAMTS-13. Blood, 2010. 115(2): p. 370–8.

40. Abdulkareem, N., et al., Proteomics in aortic aneurysm--what have we learnt so far? Proteomics Clin Appl, 2013. 7(7-8): p. 504–15.

41. Didangelos, A., et al., Proteomics characterization of extracellular space components in the human aorta. Mol Cell Proteomics, 2010. 9(9): p. 2048–62.

42. Pilop, C., et al., Proteomic analysis in aortic media of patients with Marfan syndrome reveals increased activity of calpain 2 in aortic aneurysms. Circulation, 2009. 120(11): p. 983–91.

43. van Bogerijen, G.H., et al., Biomarkers in TAA-the Holy Grail. Prog Cardiovasc Dis, 2013. 56(1): p. 109–15.

44. Ikonomidis, J.S., et al., Plasma biomarkers for distinguishing etiologic subtypes of thoracic aortic aneurysm disease. J Thorac Cardiovasc Surg, 2013. 145(5): p. 1326–33.

45. Cai, H., et al., D-Dimer Is a Diagnostic Biomarker of Abdominal Aortic Aneurysm in Patients With Peripheral Artery Disease. Front Cardiovasc Med, 2022. 9: p. 890228.

46. Vainas, T., et al., Serum C-reactive protein level is associated with abdominal aortic aneurysm size and may be produced by aneurysmal tissue. Circulation, 2003. 107(8): p. 1103–5.

47. Toumpoulis, I.K., et al., Differential expression of collagen type V and XI alpha-1 in human ascending thoracic aortic aneurysms. Ann Thorac Surg, 2009. 88(2): p. 506–13.

48. Marshall, L.M., et al., Thoracic aortic aneurysm frequency and dissection are associated with fibrillin-1 fragment concentrations in circulation. Circ Res, 2013. 113(10): p. 1159–68.

49. Syx, D., et al., Bi-allelic AEBP1 mutations in two patients with Ehlers-Danlos syndrome. Hum Mol Genet, 2019. 28(11): p. 1853–1864.

50. Thompson, R.W. and B.T. Baxter, MMP inhibition in abdominal aortic aneurysms. Rationale for a prospective randomized clinical trial. Ann N Y Acad Sci, 1999. 878: p. 159–78.

51. Xiong, W., et al., Doxycycline delays aneurysm rupture in a mouse model of Marfan syndrome. J Vasc Surg, 2008. 47(1): p. 166–72; discussion 172.

